# Cohort-HMM marker recruitment with per-OG orthology QC for phylogenomic supermatrices

**DOI:** 10.64898/2026.05.27.728348

**Authors:** Torben N. Nielsen

## Abstract

OrthoFinder’s all-vs-all DIAMOND step systematically misses single-copy orthogroups (SC OGs) at deep taxonomic divergence: a marker recovered cleanly within a tightly defined cohort is dropped when the same marker is searched against phylum-broad metagenome-assembled genome (MAG) sets, because pairwise sequence similarity falls below DIAMOND’s detection threshold even when the underlying ortholog is present. The result is biased dropout — supermatrices that retain genomes near the cohort but lose genomes from the deeper, more diverged corners of the same phylum. We describe a two-stage cohort-HMM recruitment pipeline (per-OG profile HMMs built from cohort alignments, then `hmmsearch` against the broader proteome set) followed by an independent per-OG gene-tree QC step that classifies each recruited hit relative to the cohort’s most recent common ancestor (MRCA) descendant set, with a per-MAG paralog-rate filter applied before supermatrix concatenation. We characterize the pipeline across three taxonomic ranks. At phylum scale (*Omnitrophota*, 97 cohort OGs, 714 NCBI MAGs), the recruitment recovers MAGs that the OrthoFinder-only supermatrix would otherwise drop, and the QC identifies 2 deep-peripheral MAGs — divergent genomes whose per-OG tips repeatedly place outside the cohort MRCA descendant set despite being orthologs — that the per-MAG filter removes. At family scale (*Pelagibacteraceae*, 146 cohort OGs, 366 NCBI MAGs) and at genus scale (*Actinomarina*, 289 cohort OGs, 23 NCBI MAGs), the per-tip paralog-candidate rate drops to 0.0 %. The pipeline addresses two independent failure modes. *Cohort paralog density* breaks strict-SC OG discovery at the cohort step (the family-rank case, where every candidate marker has at least one cohort species carrying multiple copies; the relaxed cohort criterion supplies the marker set and HMM recruitment disambiguates which copy each NCBI MAG contributes). *DIAMOND-reach attrition* breaks OG assignment for the most divergent NCBI MAGs (the phylum-rank case, where pairwise similarities fall below DIAMOND’s detection threshold; HMM recruitment recovers the dropouts and the per-OG QC step filters residual paralog candidates). At genus rank both modes are inactive and OrthoFinder suffices directly; HMM recruitment runs but finds no new orthologs. Code and per-case data products are released as a community resource at Zenodo (DOI 10.5281/zenodo.20422348).

## Introduction

Phylogenomic trees of bacteria and archaea are built from supermatrices: concatenated alignments of single-copy orthogroups (SC OGs) shared across the input proteomes. The orthogroup-discovery step at the front of this workflow is critical — if a marker is silently dropped or a paralog silently included, the resulting tree carries a bias that no downstream diagnostic will catch. OrthoFinder (Emms et al. 2025) is the de facto tool for this step, and is reliable when applied to a *tight* input cohort — a genus, a family, or a small clade where pairwise sequence similarity between every pair of genomes stays above DIAMOND’s detection threshold.

At wider taxonomic scope, the same workflow fails silently. When the input set extends to all GTDB representatives of a phylum — species-representative metagenome-assembled genomes selected via GTDB and downloaded from NCBI GenBank (GCA accessions) or RefSeq (GCF accessions), hereafter the *NCBI pool* — pairwise similarities for the most divergent lineages fall below DIAMOND’s detection threshold, and OrthoFinder’s DIAMOND step fails to identify the orthologs that link those lineages to the rest of the input. OrthoFinder still runs to completion, still reports a clean per-cohort SC OG count, still builds a supermatrix; the supermatrix just systematically drops the deepest-branching MAGs at the gap-coverage cutoff. In the *Omnitrophota* case study below, the OrthoFinder + DIAMOND baseline drops 175 of 710 species representatives at a ≥ 90 % gap cutoff against a 97-OG cohort supermatrix. The dropped MAGs are not low-quality: median CheckM2 completeness 94 %, contig count uncorrelated with the supermatrix gap rate (Spearman ρ = 0.03). They are exactly the lineages whose phylogenetic placement matters most.

We address this by replacing the pairwise-similarity step with profile-based detection on the NCBI side, while keeping OrthoFinder’s marker definition on the cohort side. Profile HMMs (Eddy 2011) are built from the per-OG cohort alignments — where the marker definition is unambiguous — and used to identify the corresponding ortholog in each NCBI proteome via `hmmsearch`. An independent per-OG gene-tree quality control step then asks whether each recruited NCBI tip places inside the descendant set of the cohort MRCA on the per-OG gene tree (ortholog) or outside (paralog candidate); a per-MAG paralog-rate filter excludes MAGs that the gene trees flag at a rate above threshold. The HMMs supply detection power that DIAMOND cannot reach; the gene-tree QC supplies the orthology call on an independent signal.

The pipeline acts on two independent failure modes. *Cohort paralog density* is a property of the cohort alone: when paralog-bearing species accumulate, OrthoFinder’s strict-SC criterion can fail to yield any usable OGs. *DIAMOND-reach attrition* is a property of the NCBI pool relative to the cohort: deeply divergent NCBI MAGs fall below DIAMOND’s detection threshold and lose OG membership. We characterize the pipeline against three case studies chosen to span this space: *Actinomarina* at genus rank, where neither failure mode is active; *Pelagibacteraceae* at family rank, where cohort paralog density is active; and *Omnitrophota* at phylum rank, where DIAMOND-reach attrition is active. The three cases show when each step of the pipeline contributes and when the pipeline runs without changing the OrthoFinder-only result.

## Methods

### Overview

For each case study, the pipeline is:

1. **Cohort definition**. Pick a tightly defined cohort of genomes at the desired taxonomic scope. The cohort should be cohesive enough that OrthoFinder’s DIAMOND step works on it directly.
2. **Cohort SC OG discovery**. Run OrthoFinder on cohort proteomes only; we consume only `Orthogroups.txt` (orthogroup membership) downstream and discard or terminate the per-OG MSA and gene-tree phase. Extract single-copy or relaxed single-copy OGs from that file. We use OrthoFinder v3.1.4 (Emms et al. 2025) with DIAMOND backend (Buchfink et al. 2021), 32–64 threads per case.
3. **HMM build**. For each SC OG, build a profile HMM from the cohort MAFFT L-INS-i alignment (`--localpair --maxiterate 1000`, with `--anysymbol` for the *Pelagibacteraceae* and *Actinomarina* cases; Katoh & Standley 2013) via `hmmbuild` (HMMER 3.4; Eddy 2011). Concatenate all per-OG HMMs and run `hmmpress`.
4. **Broader proteome fetch**. Select the GTDB R232 species representatives (GTDB-Tk R232; Chaumeil et al. 2022) matching the taxonomic scope and download the corresponding assemblies from NCBI GenBank (GCA accessions) or RefSeq (GCF accessions). Call genes locally with Pyrodigal v3.6.3 (`force_nonsd=True`; Larralde 2022) rather than relying on NCBI annotation, which is patchy for GenBank-only deposits.
5. **HMM scan**. Run `hmmsearch` of the cohort HMM panel against the concatenated NCBI-pool proteomes. For each NCBI genome and each OG, retain the best hit by bit score.
6. **Per-OG gene-tree QC**. For each OG, write a per-OG FASTA combining the cohort sequences plus the single best NCBI hit per genome, align with MAFFT L-INS-i (`--localpair --maxiterate 1000`), and infer a per-OG tree under LG+G4 with `-fast` — the per-OG QC tree, not the supermatrix tree of step 9. Identify the most recent common ancestor (MRCA) of the cohort tips (excluding any baked-in outgroups) and classify each NCBI tip as ortholog (inside the cohort MRCA descendant set) or paralog candidate (outside).
7. **MAG filter**. For each NCBI MAG, compute the per-MAG paralog-candidate rate across the OGs at which the MAG had an HMM hit. MAGs with a per-MAG rate > 25 % are dropped from the supermatrix. Per-(MAG, OG) cells that are paralog candidates are also dropped, even when the MAG itself passes the rate filter.
8. **Supermatrix**. For each retained OG, take the per-OG QC alignment (MAFFT L-INS-i, built in step 6), filter to the cohort + retained NCBI ortholog set, and trim with BMGE2 (`-t AA -m BLOSUM30 -e 0.5 -g 0.5 -b 3`; Criscuolo & Gribaldo 2010); concatenate into a partitioned supermatrix.
9. **Tree inference**. Infer the supermatrix tree with IQ-TREE v3.0.1 (Wong et al. 2026) under `LG+F+R10 -B 1000 -alrt 1000 -bnni`, 32 threads, seed 42. Outgroups, when available in the cohort, are passed via `-o`; otherwise the tree is rerooted at midpoint post-hoc.

### Cohort SC-OG criterion: strict vs relaxed

At genus rank (*Actinomarina* case study), the cohort yields strict single-copy OGs — present in 100 % of cohort species, with no paralog copies. At family rank (*Pelagibacteraceae*), the strict criterion yields 0 OGs across 539 cohort species, because at least one paralog copy appears in some species for every candidate marker; relaxing to ≥ 95 % cohort presence and ≤ 1 % paralog-bearing species yields a workable cohort marker set. At phylum rank (*Omnitrophota*), 97 strict single-copy OGs are recovered from a 184-proteome discovery cohort drawn from Nielsen (in preparation) — complete in-house genomes plus outgroup tips and four named taxonomic anchors. The 97-OG HMM panel is then applied to the 229-genome supermatrix cohort (176 complete + 53 high-quality in-house genomes) and to the 714-MAG GTDB R232 species-representative pool downstream. The strict criterion still works at this rank because the discovery cohort itself is constrained, not the broader MAG set. The strict vs relaxed choice is therefore made at the cohort step, not by the recruitment.

### Per-OG QC implementation

`qc_03_classify_orthology.py` reads the per-OG gene tree built in step 6 and uses ete4 (`Tree`, `common_ancestor`; Huerta-Cepas, Serra & Bork 2016) to find the MRCA of the cohort tips, excluding any tips that were defined as outgroups in the cohort. For each non-cohort tip in the tree, the classifier reports `ortholog` if the tip falls within the descendants of the cohort MRCA and `paralog_candidate` otherwise.

The `paralog_candidate` label has known biological ambiguity: it collapses three cases — a true paralog (the HMM recruited the wrong copy), a deep peripheral ortholog (the gene is orthologous but its per-OG topology places the tip outside the cohort MRCA descendant set by genuine divergence), and a long-branch artifact (limited per-OG signal places the tip incorrectly). At phylum rank these three cases are not separable from a single per-OG tree.

The per-MAG paralog-rate filter (step 7) addresses this ambiguity in aggregate. A MAG whose per-OG QC consistently flags it as outside the cohort MRCA descendant set across many OGs (≥ 25 % of OGs) is read as a deep peripheral position rather than per-OG noise. The filter does not distinguish among the three cases above, but removes MAGs that all three predict will introduce supermatrix bias.

### Outgroup convention

The genus-rank (*Actinomarina*) and family-rank (*Pelagibacteraceae*) supermatrix trees are inferred with no explicit `-o` outgroup flag and rooted post-hoc; this mirrors the convention used in the *Pelagibacter* family-rank paper (Nielsen & Lui 2026a). The phylum-rank (*Omnitrophota*) tree uses an explicit four-tip Planctomycetota–Verrucomicrobiota–Chlamydiota (PVC) clade outgroup at inference time — four in-house contigs (two from SFE sample 1W, one from 8W, one from B7), listed in the per-case IQ-TREE log — because those outgroup tips were part of the original cohort definition and are needed to root the topology at the *Omnitrophota* / sister-PVC split.

### Implementation

Per-case directories share a uniform layout: cohort and NCBI-pool proteomes (`input/`), profile HMMs (`hmms/`), raw `hmmsearch` output (`hmmsearch_tbl/`), per-OG alignments and QC trees (`per_og_aln/`, `per_og_tree/`), the concatenated and partitioned supermatrix (`supermatrix_hmm/`), and the final IQ-TREE outputs (`iqtree_hmm/`). The pipeline scripts mirror across the two cases run in this paper (*Pelagibacteraceae, Actinomarina*); the *Omnitrophota* case reuses the discovery cohort and per-OG alignments from Nielsen (in preparation) and starts from the HMM build. The full case-study data products — recovery TSVs, per-OG QC tables, alignments, HMM panels, supermatrices, and IQ-TREE outputs — are released at Zenodo (DOI 10.5281/zenodo.20422348).

## Results

### Case study 1 — *Omnitrophota* (phylum rank): recruitment and QC are both load-bearing

Cohort SC OG discovery was carried out on the 184-proteome *Omnitrophota* discovery cohort (complete in-house genomes plus outgroup tips and four named taxonomic anchors; Nielsen, in preparation), yielding 97 strict single-copy orthogroups under default OrthoFinder. The 97-OG HMM panel was then applied to the 229-genome supermatrix cohort (176 complete + 53 high-quality in-house genomes) and to the 714-MAG GTDB R232 *Omnitrophota* species-representative pool. Scanning recovered a median of 97 OGs per NCBI MAG (the full panel), with the lowest-coverage MAGs at roughly 50 OGs. Without the recruitment, the OrthoFinder + DIAMOND baseline on the same NCBI pool drops 175 of 710 MAGs at a ≥ 90 % gap cutoff against the 97-OG cohort supermatrix: the dropouts the recruitment recovers.

The per-OG gene-tree QC then placed 72,092 of 73,510 recruited tip placements inside the cohort MRCA descendant set (98.1 %; 1.9 % paralog candidates). The in-house high-quality fraction returned 0.1 % paralog candidates (6 / 5,104 placements); the NCBI fraction returned 2.1 % (1,412 / 68,406). Aggregating to the per-MAG level, two MAGs exceeded the 25 % per-MAG paralog-rate threshold and were excluded from the supermatrix; an additional 7 NCBI MAGs were dropped at the per-OG gap-coverage step at supermatrix build. With cohort + outgroups + retained NCBI recruited tips, the IQ-TREE input is 938 taxa × 23,018 amino acid sites (229 in-house cohort + 4 PVC outgroups + 705 NCBI MAGs retained from the 714-MAG R232 pool); the inferred tree has log-likelihood −11,440,270.3 (consensus −11,440,200.4; 1,000-rep ultrafast bootstrap, UFBoot; Figure 4).

The *Omnitrophota* case is the canonical failure case for the cohort-fit problem: it has 175 OrthoFinder + DIAMOND dropouts to recover and a 1.9 % per-tip paralog-candidate rate (> 25 % at the MAG level for 2 deep-peripheral MAGs) at the same time. Both recruitment and QC contribute.

### Case study 2 — *Pelagibacteraceae* (family rank): recruitment is essential, QC returns no flags

A 539-proteome cohort drawn from the *Pelagibacter* family paper’s collection (Nielsen & Lui 2026a) yielded 0 strict single-copy OGs under default OrthoFinder: every candidate marker had at least one cohort species carrying multiple copies. Relaxing to ≥ 95 % cohort presence and ≤ 1 % paralog-bearing species yielded 146 cohort marker OGs. Building per-OG HMMs from this cohort and scanning 366 GTDB R232 species representatives recovered a median of 141 OGs per MAG (mean 139.5 ± 6.8; 366/366 MAGs passed the < 90 % gap cutoff, 327/366 passed < 10 % gap). The HMM recruitment was therefore the only path to a usable marker set at this cohort scale: the strict-SC criterion failed at the cohort step, and without the relaxed cohort + HMM recruitment there would have been no supermatrix to build.

The per-OG gene-tree QC placed 50,789 of 50,789 NCBI tip placements inside the cohort MRCA descendant set (100.0 %; 0 paralog candidates across all 146 OGs). No MAGs were dropped on the per-MAG paralog rate. The QC step ran, returned a uniformly clean result, and confirmed that the recruitment did not manufacture spurious orthologs — but the same supermatrix would have resulted from the recruitment alone. With cohort + NCBI MAGs, the IQ-TREE input is 905 taxa × 20,436 amino acid sites; the inferred tree has log-likelihood −3,365,680.1 (consensus −3,365,329.1; 1,000-rep UFBoot; Figure 2).

**Figure 1.**
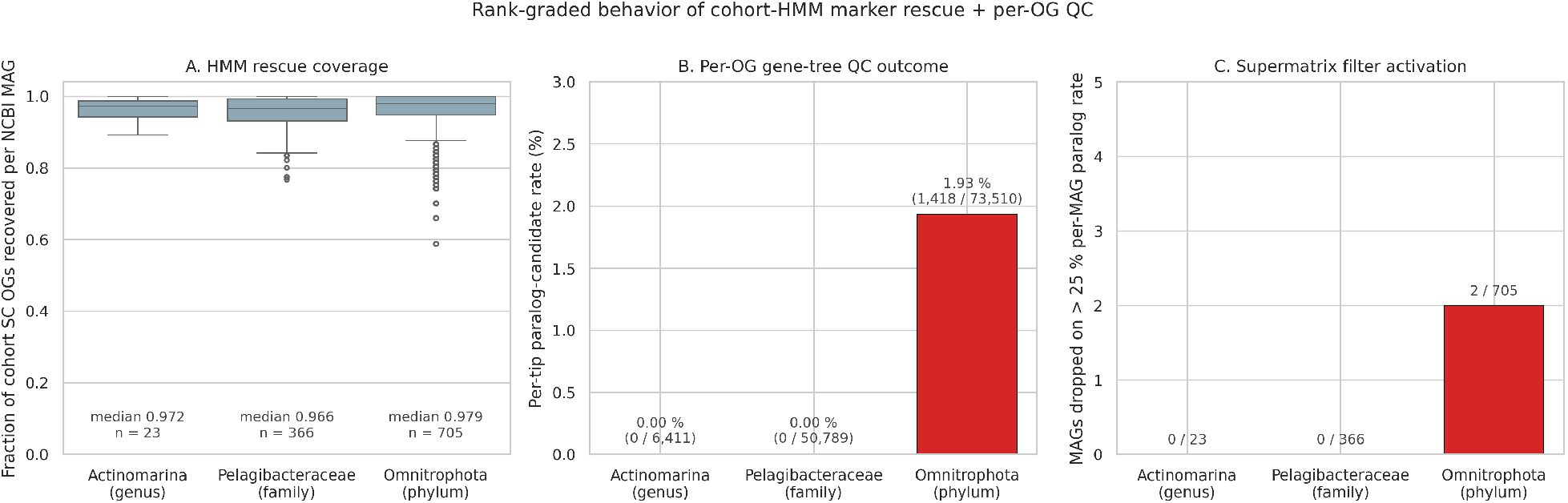
Recruitment-rate and paralog-rate by cohort rank. Three panels across the case studies: (A) median OGs recovered per NCBI MAG by HMM recruitment, normalized to cohort SC OG count; (B) per-tip paralog-candidate rate after per-OG gene-tree QC; (C) number of MAGs dropped on the > 25 % per-MAG paralog-rate filter. The rank-graded pattern is evident: per-MAG OG recovery via HMM recruitment is uniformly high (97 %+ at all three ranks), the paralog-candidate rate scales with cohort divergence (0.0 % / 0.0 % / 1.9 % for *Actinomarina* / *Pelagibacteraceae* / *Omnitrophota*), and the per-MAG filter activates only at phylum rank.

**Figure 2.**
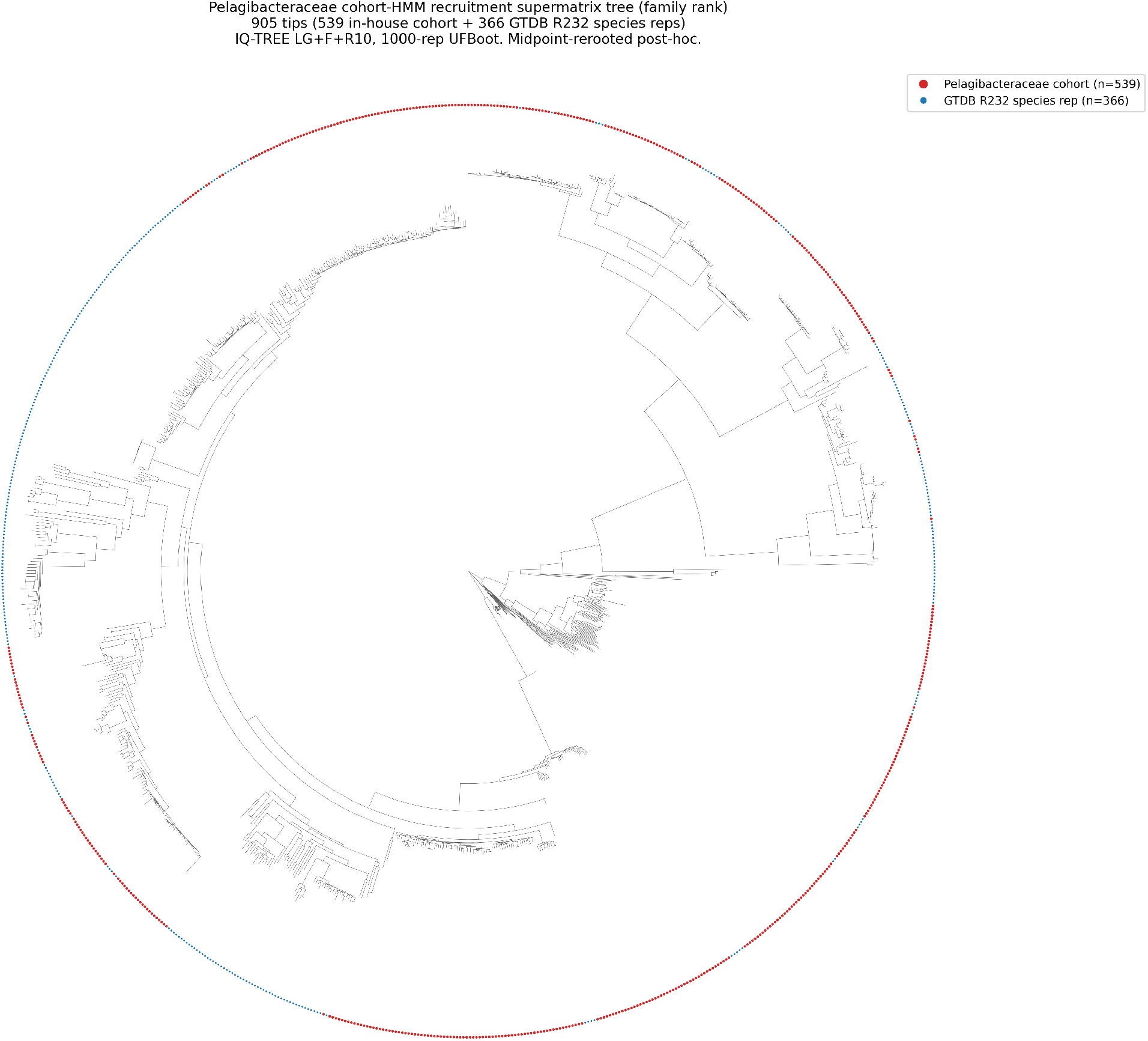
*Pelagibacteraceae* cohort-HMM recruitment supermatrix tree (905 tips). Family-rank worked example. 539 in-house *Pelagibacteraceae* cohort tips (red, from Nielsen & Lui 2026a) and 366 GTDB R232 species representatives (grey) recovered by HMM recruitment. Per-OG gene-tree QC returned 0 paralog candidates across 50,789 placements, so all 366 NCBI MAGs are retained. Inferred under IQ-TREE v3.0.1 LG+F+R10, 1,000-rep UFBoot, no outgroup specified; rerooted at midpoint post-hoc for display.

The *Pelagibacter* case shows that at family rank the recruitment is essential but the QC returns no flags — the family-rank cohort’s gene-tree placement is dense enough that the recruited NCBI tips fall reliably inside the cohort MRCA descendant set without needing to be tested.

### Case study 3 — *Actinomarina* (genus rank): the recruitment is not needed

A 144-proteome cohort drawn from the *Actinomarina* genus paper’s collection (Nielsen & Lui 2026b) yielded 289 strict single-copy OGs under default OrthoFinder. Building per-OG HMMs from this cohort and scanning the 23 GTDB R232 species representatives recovered a median of 281 OGs per MAG (mean 278.7 ± 11.4; 23/23 MAGs passed the < 90 % gap cutoff, 22/23 passed < 10 %).

The per-OG gene-tree QC placed 6,411 of 6,411 NCBI tip placements inside the cohort MRCA descendant set (100.0 %; 0 paralog candidates across all 289 OGs). With cohort + NCBI MAGs, the IQ-TREE input is 167 taxa × 77,860 amino acid sites; the inferred tree has log-likelihood −1,729,333.5 (consensus −1,729,336.8; 1,000-rep UFBoot; Figure 3).

**Figure 3.**
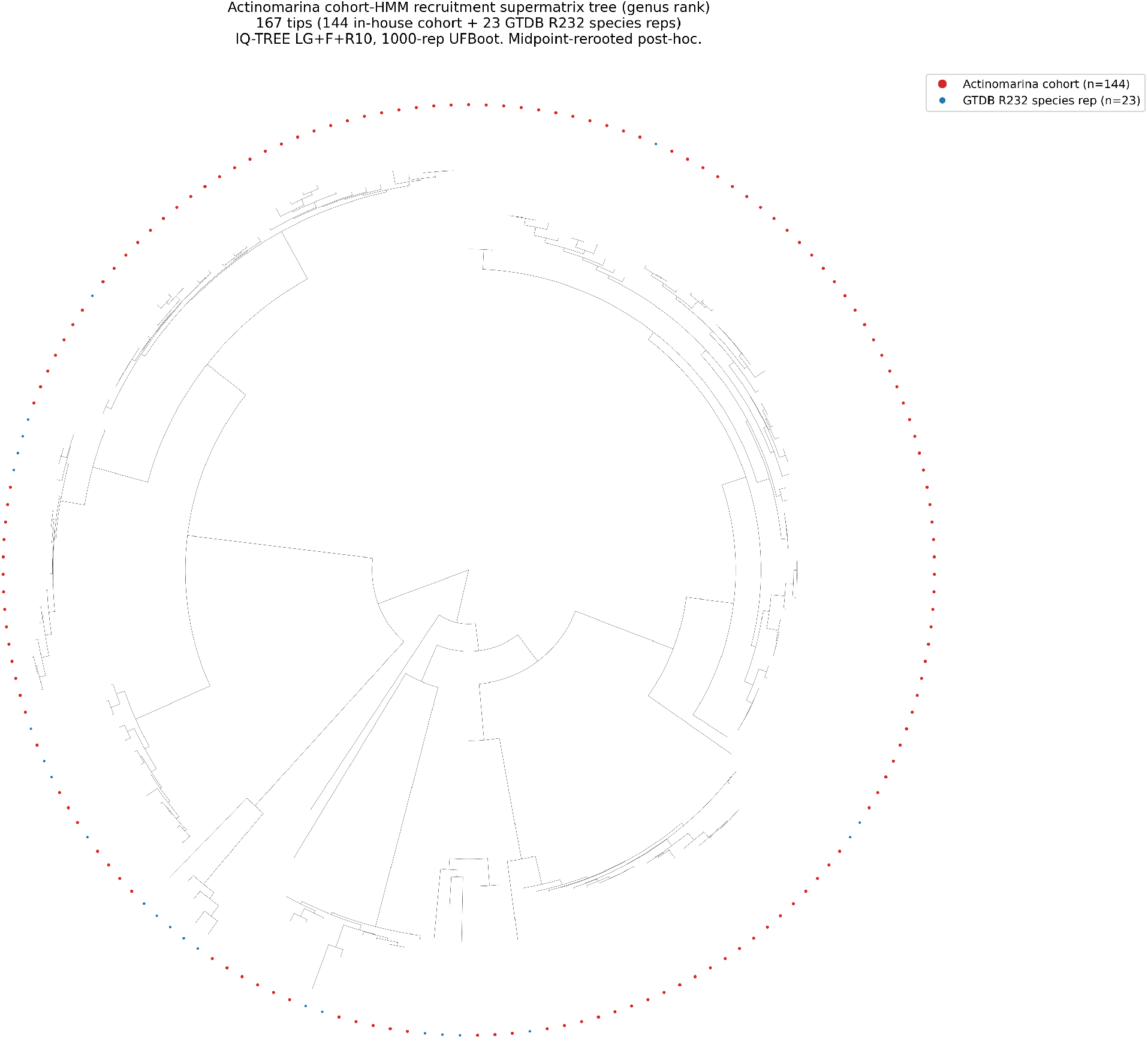
*Actinomarina* cohort-HMM recruitment supermatrix tree (167 tips). Genus-rank worked example. 144 in-house *Actinomarina* cohort tips (red, from Nielsen & Lui 2026b) and 23 GTDB R232 species representatives (grey) recovered by HMM recruitment. Per-OG gene-tree QC returned 0 paralog candidates across 6,411 placements. Inferred under IQ-TREE v3.0.1 LG+F+R10, 1,000-rep UFBoot, no outgroup specified; rerooted at midpoint post-hoc for display.

**Figure 4.**
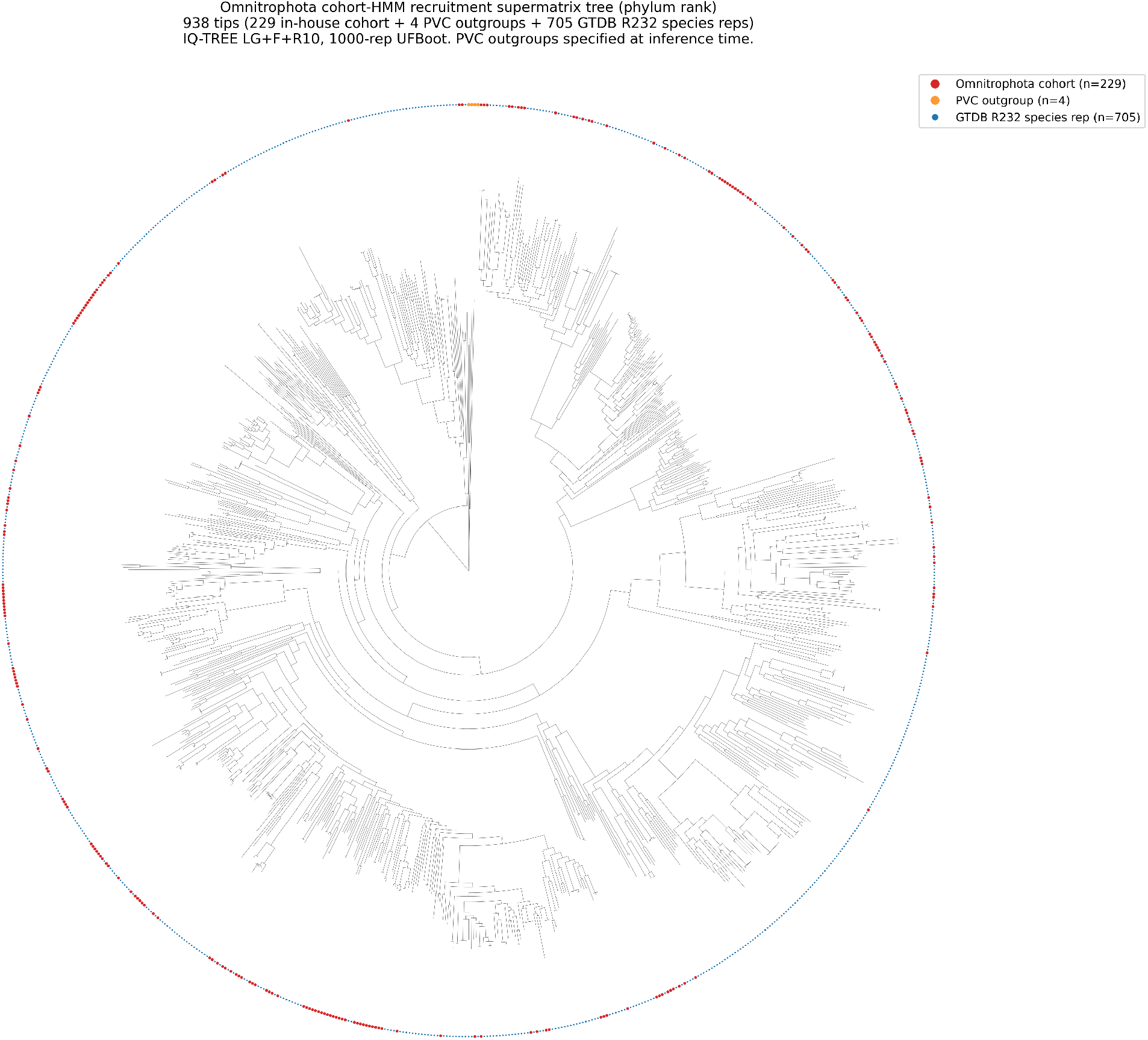
*Omnitrophota* cohort-HMM recruitment supermatrix tree (938 tips). Phylum-rank worked example. 229 in-house *Omnitrophota* cohort tips (red; 176 complete + 53 high-quality genomes from Nielsen, in preparation), 4 PVC outgroup tips (orange), and 705 GTDB R232 species representatives (grey) recovered by HMM recruitment and retained after per-OG and per-MAG filtering (from a 714-MAG R232 pool; 2 MAGs dropped at > 25 % per-MAG paralog rate, 7 dropped at the per-OG gap-coverage step). Inferred under IQ-TREE v3.0.1 LG+F+R10, 1,000-rep UFBoot, with PVC outgroups specified at inference time. This tree is used as the phylogenomic backbone of the *Omnitrophota* paper (Nielsen, in preparation).

The *Actinomarina* case is the negative control. At genus rank, the original OrthoFinder DIAMOND step recovered the markers without needing recruitment; the HMM recruitment and the per-OG QC both ran cleanly but returned no actionable signal beyond what OrthoFinder already produced. The case is included here to show that the pipeline does not introduce spurious recruitments or paralog flags at the shallow end — both stages return no flags when the cohort fit is already tight.

### Recruitment rate vs cohort scale

Across the three cases, the per-tip paralog-candidate rate scales with cohort divergence (Figure 1): 0.0 % at genus rank (*Actinomarina*), 0.0 % at family rank (*Pelagibacteraceae*), 1.9 % at phylum rank (*Omnitrophota*). The HMM-recruitment recovery rate likewise scales: at genus rank the median MAG recovers 281/289 OGs (97 %) where the original DIAMOND step already worked; at family rank 141/146 (97 %) where the relaxed cohort criterion was the bottleneck; at phylum rank 175 MAGs are recovered that the cohort + DIAMOND baseline would have dropped at the ≥ 90 % gap cutoff, with the per-MAG QC filter then removing 2 deep-peripheral MAGs from the recruited pool (705 of 714 retained; the remaining 7 are dropped at the per-OG gap-coverage step).

## Discussion

### When to use the recruitment

Two independent diagnostics determine whether HMM recruitment is needed, and they target different failure modes.

*Cohort paralog density* is diagnosed at the cohort step. If OrthoFinder’s strict-SC criterion yields zero (or few) OGs on the cohort proteomes alone, the cohort itself carries enough paralog-bearing species that strict orthology cannot converge. The relaxed criterion (≥ 95 % presence, ≤ 1 % paralog-bearing species) recovers a marker set, and HMM recruitment is then needed to disambiguate which copy of each multi-copy OG each NCBI MAG should contribute to the supermatrix. This is the *Pelagibacteraceae* family-rank case (0 strict-SC OGs, 146 relaxed-cohort OGs).

*DIAMOND-reach attrition* is diagnosed at the NCBI-pool step. The signal is the number of NCBI MAGs dropped at a ≥ 90 % gap cutoff against the OrthoFinder + DIAMOND cohort + NCBI supermatrix. If the answer is zero or near-zero, DIAMOND is reaching the NCBI pool from the cohort and HMM recruitment adds nothing on the NCBI side. If the answer is substantial — 175 of 710 in the *Omnitrophota* phylum-rank case — pairwise similarities for the deepest-branching MAGs have fallen below DIAMOND’s detection threshold and HMM recruitment is needed to recover them via profile-based detection. The per-OG QC step is needed alongside the recruitment because deep peripheral lineages, true paralogs, and long-branch artifacts all surface at this scale.

The two diagnostics are orthogonal: cohort paralog density is a property of the cohort itself, while DIAMOND-reach attrition is a property of the NCBI pool relative to the cohort. The *Actinomarina* genus-rank case has neither — strict-SC succeeds, DIAMOND reaches all NCBI species reps, HMM recruitment runs and finds no new orthologs.

### Limitations

The per-OG QC step has three limitations. First, the `paralog_candidate` classification at phylum rank collapses three cases — true paralogs, deep peripheral orthologs, and long-branch artifacts — which are not separable from a single per-OG gene tree. The per-MAG rate filter aggregates over this ambiguity and is calibrated to identify MAGs whose paralog rate is high enough to plausibly affect supermatrix placement, but the filter applies uniformly across all three without distinguishing them. A reader interested in any single MAG’s classification should treat the label as a placement flag, not a paralog claim.

Second, the cohort MRCA-based classifier depends on the cohort being phylogenetically cohesive. When the cohort spans deep within-cohort divergences — for example, the *Gorgyraeia* / *Omnitrophia* split within *Omnitrophota* — the MRCA includes a larger fraction of the tree and the classifier is more permissive than at a tight cohort such as *Actinomarina*. The 25 % per-MAG threshold is calibrated against the *Omnitrophota* case-study distribution; tighter cohorts will have lower null rates and may warrant a tighter threshold.

Third, the HMM recruitment is conservative against false negatives but not against false positives. An HMM built from a cohort alignment will recruit the best match in the broader proteome to that HMM, whether or not the broader proteome carries a true ortholog — if a paralog is the closest match in protein space, the pipeline will accept it. The per-OG QC step flags this case, and the per-MAG filter catches the systematic case where it recurs across many OGs in the same MAG.

### Relationship to other tools

GTDB-Tk’s `classify_wf` (Chaumeil et al. 2022) uses pplacer placement onto the GTDB reference tree’s bac120 / ar53 marker concatenation, and is not subject to the dropout failure mode described here because pplacer does HMM-based marker calling rather than pairwise similarity. Per-genome classification is not affected by the cohort-fit problem. The cohort-fit problem appears specifically in *cohort-derived* supermatrix inference, where the marker set is defined by the cohort itself rather than by a fixed reference HMM panel like bac120. Within that scope, the failure mode is well-characterized for OrthoFinder + DIAMOND (Emms et al. 2025) and the recruitment is targeted at that specific failure mode.

## Conclusion

The standard OrthoFinder + DIAMOND supermatrix workflow is vulnerable to two silent failure modes that the workflow itself does not detect: cohort paralog density at the cohort step, and DIAMOND-reach attrition at the NCBI side. Cohort-HMM recruitment with per-OG orthology QC addresses both, replacing pairwise similarity with profile detection where DIAMOND cannot reach while preserving OrthoFinder’s marker definition on the cohort side, and adding a per-OG gene-tree QC step that calls orthology on a signal independent of the recruitment. The three case studies show the pipeline behaviour is rank-graded — both steps are essential at phylum rank, the recruitment alone at family rank, neither at genus rank — and the two diagnostics in *When to use* let a user decide whether the pipeline is needed for a given cohort plus NCBI pool combination. The pipeline is reproducible from input proteomes alone with no manual curation.

## Cross-references

The *Pelagibacteraceae* family-rank case study uses the published *Pelagibacter* genome collection of Nielsen & Lui (2026a). The *Actinomarina* genus-rank case study uses the *Actinomarina* genome collection (Nielsen & Lui 2026b). The *Omnitrophota* phylum-rank case study shares its phylogenomic backbone with the *Omnitrophota* genome paper (Nielsen, in preparation), which uses the cohort-HMM recruitment tree directly (Figure 2 of that paper); the present paper’s *Omnitrophota* case study reports the method by which that tree was inferred.

## Notes

### Competing Interest Statement

The authors have declared no competing interest.

